# Coexistence of long and short DNA constructs within adhesion plaques

**DOI:** 10.1101/2020.05.12.090357

**Authors:** Long Li, Mohammad Arif Kamal, Henning Stumpf, Franck Thibaudau, Kheya Sengupta, Ana-Sunčana Smith

**Author notes:** contributed equally (Long Li, Mohammad Arif Kamal).

## Abstract

Adhesion domains forming at the membrane interfaces between two cells or a cell and the extracellular matrix commonly involve multiple proteins bridges. However, the physical mechanisms governing the domain structures are not yet fully resolved. Here we present a joint experimental and theoretical study of a mimetic model-system, based on giant unilammelar vesicles interacting with supported lipid bilayers, with which the underlying physical effects can be clearly identified. In our case, adhesion is induced by simultaneous action of DNA linkers with two different lengths. We study the organization of bridges into domains as a function of relative fraction of long and short DNA constructs. Irrespective of the composition, we systematically find adhesion domains with coexisting DNA bridge types, despite their relative differences in length of 9 nm. However, at short length scales, below the optical resolution of the microscope, simulations suggest the formation of nanodomains by the minority fraction. The nano-aggregation is more significant for long bridges, which are also more stable, even though the enthalpy of membrane insertion is the same for both species.

---

Cell adhesion is a fundamental biological process that relies on the formation of complex macromolecular assemblies, involving a number of proteins with different binding affinities, elastic properties and lengths ^1^. These proteins may mediate adhesion in either a sequential manner, as it is the case of weak selectin adhesion preceding strong integrin adhesion in rolling leukocytes ^2–4^, or simultaneously, for example during the formation of immune synapses. In this case T-cell receptors and integrins segregate ^5,6^, presumably because of the differences in the length of their respective adhesive constructs ^7^. However, the full understanding of the length-induced separation is still missing.

One proven approach to study these mechanisms is using cell mimetic systems consisting of functionalized giant unilamelar vesicles (GUVs) adhering to supported lipid bilayers (SLB). So far, most mimetic adhesion systems involved only one protein pair, or mixtures between binders and repellers ^8,9^, while the design of systems consisting of more than one binding pair is still challenging, and an active field of studies ^10^.

Constructing theoretical models from first principles and understanding the interplay between multiple binding pairs during adhesion has been equally demanding. The difficulty here is to accurately capture membrane-mediated interactions over the large span of time and length scales involved in the nucleation and growth of adhesions. Over the last decades, the evolution of large domains with the two binding pairs fully separated were discussed extensively ^7,11,12^. More recently, small stable domains, as well as adhesion structures containing more than one type of bridge were reported in simulations ^13^, but to this date, these results did not receive experimental confirmation.

Here we present a synergistic experimental and theoretical effort to address this regime. Building on our previous work on mimetic systems ^9^, we functionalize the GUVs and the SLB with a binary mixture of DNA constructs that can bridge the two membranes and mediate adhesion. Such DNA constructs are particularly attractive because their length, flexibility and binding strength can be readily modified ^14–21^. We support these experimental advances by extending the recently developed Monte Carlo scheme for modeling adhesion with multiple types of ligand-receptor pairs ^13^, to modeling the formation of DNA bridges of different lengths. We systematically change the relative concentration of short and long DNA constructs and observe a continuous change in the average height of a densely packed adhesion domain both experimentally and in simulations with no clear self-segregation. This shows that adhesion structures with bridges of a significantly different length may coexist in equilibrium.

For our experiments, we prepared GUVs (98 mol% SOPC and 2 mol % DSPE-PEG2000) using the well established electroswelling method in a glucose solution, as described previously ^21^. Subsequently, vesicles were deflated and functionalized by immersion into a PBS buffer containing 6 nmol binary mixture of short and long DNA linkers, with a 40 mOsm higher osmolarity than the swelling solution.

DNA-decorated vesicles were injected into the measuring chamber and allowed to sediment to the SLB. The latter were prepared with a film b alance (Nima, C oventry, U K) applying the Langmuir-Blodgett Langmuir-Schäfer technique ^22^, at transfer pressure of 20 mN/m. SLBs consisted of pure SOPC in the proximal layer, while the distal layer facing the vesicles was formed by SOPC with 2 mol % DSPE-PEG 2000 and 1 mol % NBD-PE. The SLB as well as the GUV get decorated by DNA linkers upon the insertion of the DNA containing buffer.

The linkers were constructed form ss-DNA sequences (*N* = 20?33?(needs checking) base pairs out of which 10 base pairs build a 3 nm long sticky end, while the backbone terminates with the 4 base pair spacer attached to a cholesterol moiety). The identical segments are allowed to recombine at the sticky end to make a strand twice as long with a cholesterol at each end. Furthermore, 17 and 30 base pair antisense sequences of the backbone are allowed to recombine into respective short and long ds-DNA linkers. Because of the strong affinity of cholesterol for the lipid membrane, the DNA linker spontaneously adsorbs on the SLB and the GUV. Adhesion happens when the linker translocates one of the cholesterol moieties from the GUV to the SLB membrane or vice versa, or if it adsorbs directly from the solution with one cholesterol moiety in each membrane.

The adhesion zone between the GUV and the SLB is imaged using Reflection Interferance Contrast Microscopy (RICM), following the usual practice ^9^. Using RICM, it is possible to determine the membrane height with about 4 nm resolution. As always, to secure the correct interpretation of height (Fig. 2a), the refractive indices of the vesicle solution and the outer buffer were measured for each experiment with an Abbé Refractometer (Kruss, Germany). No adhesion is observed in the control sample when DNA-linkers are not introduced.

**Fig. 1.**
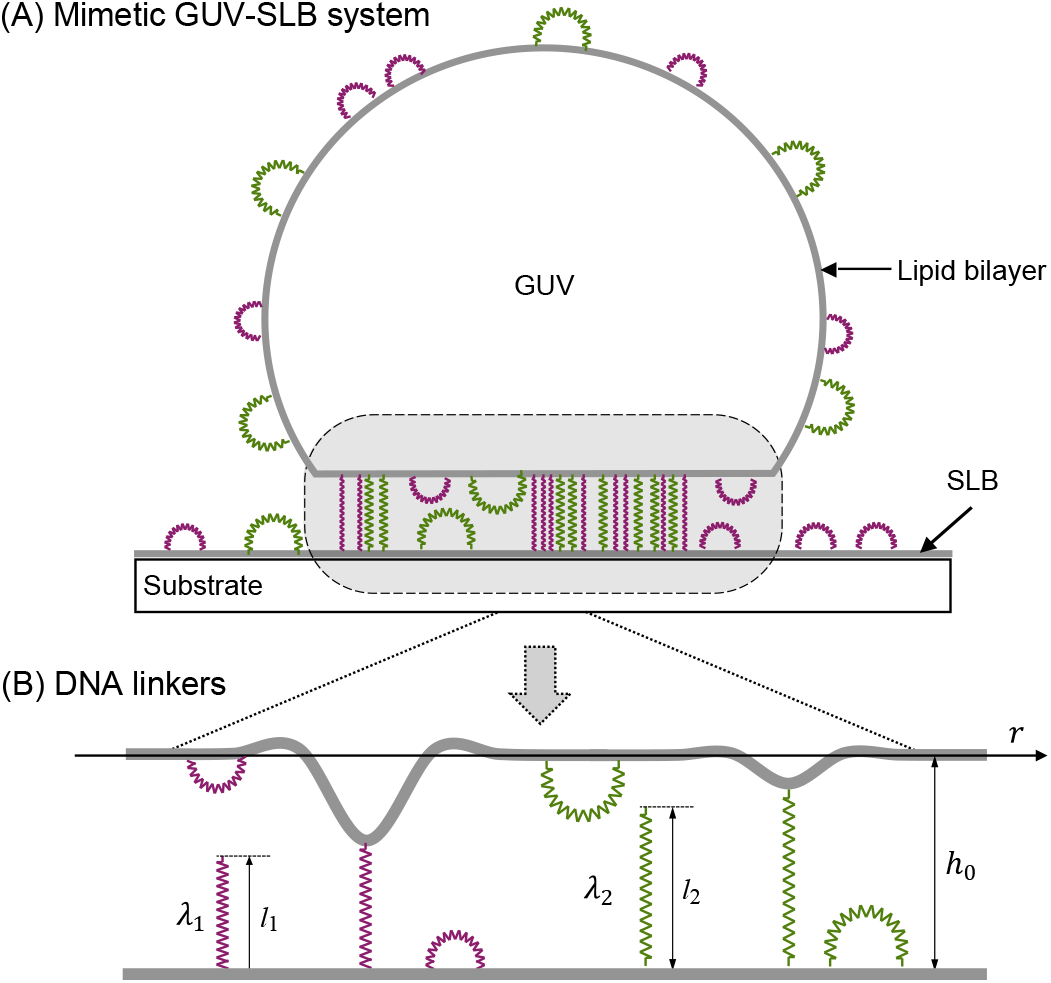
(A) GUV and SLB functionalized with ligands and receptors, respectively. When GUV approaches the a contact zone with the SLB forms, where the membranes, in the absence of bridges adopt a distance *h*_0_ of 50-100 nanometers due to the presence of a weak potential produced by various nonspecific interactions. (B) Model of a flexible membrane (thick line) hovering in a non-specific potential over a flat and rigid adherent surface. DNA constructs (B) of a length *l_i_* are modelled as elastic springs of stiffness *λ_i_*. When a DNA linker swaps from an intra-membrane to an inter-membrane configuration, a bridge is formed. Consequently, the GUV membrane deforms adopting a mean height 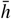, while its fluctuation amplitude *σ* is suppressed.

**Fig. 2.**
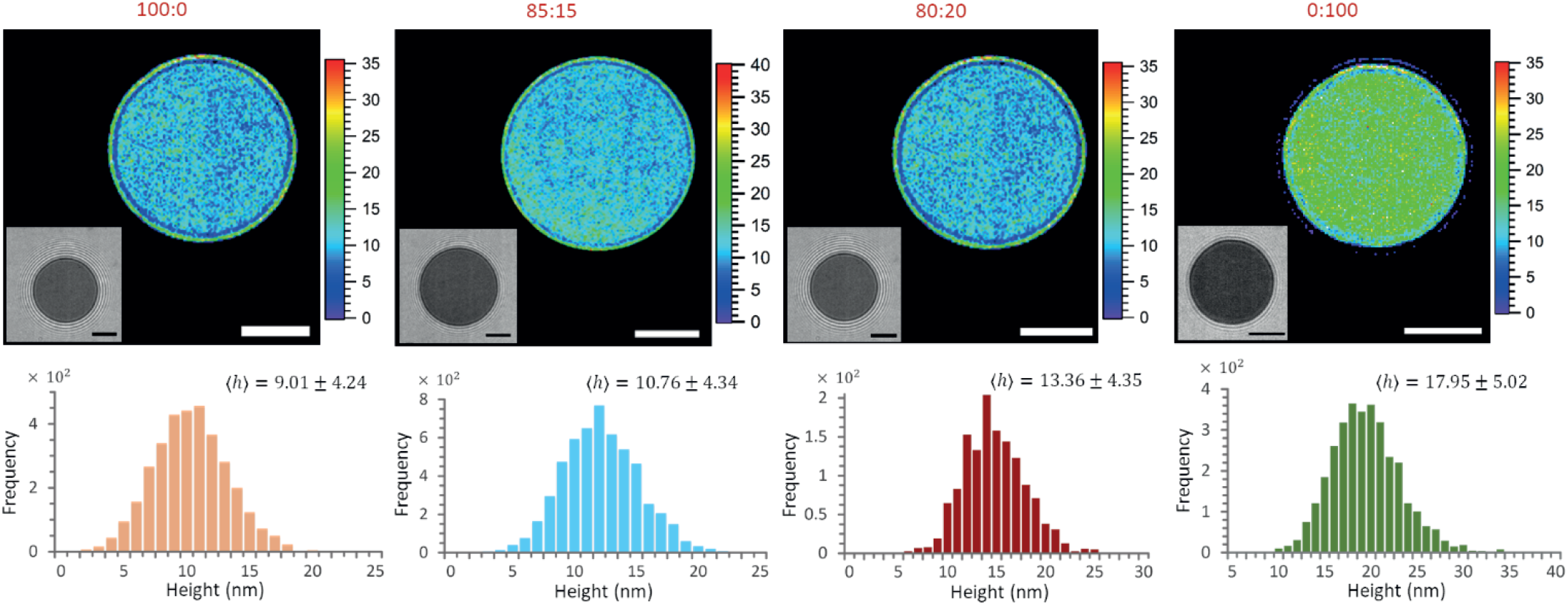
GUV-SLB adhesion mediated a binary mixture of long and short DNA ds-DNA (8nm:36nm) at fixed concentration of 6nM. Representative RICM image (inset) of the adhesion zone between the GUVs and the SLBs, corresponding height map together with height histograms at different compositions of the binary mixtures of ds-DNA. Scale bar: 5 *μ*m.

Irrespective of the DNA linkers presented on the GUV and the SLB, vesicles always adopt a uniform contact zone (insets in Fig. 2), with a roughness of about 4 nm, which is comparable to the background noise at this intensity. This small roughness suggests that the linkers are relatively densely distributed in the contact zone. Nonetheless, the dynamics of growth, as well as systematically discoid contact areas between GUVs and the SLB suggest that the linkers, even in bridge configuration, maintain certain mobility, which indicates that close packing of DNA linkers is not achieved. However, no change in the pattern is observed even several hours upon bringing GUVs into contact with the SLB, which is consistent with no ongoing restructuring, and the full equilibration of the system.

After averaging over a large number of vesicles (72 and 51 for short and long DNA-linkers, respectively), we find the GUV-SLB separation of about 9 and 18 nm, when only short or only long DNA linkers are present. However, as a binary mixture is introduced in the ratio of short:long=85:15 (58 vesicles), or 80:20 (72 vesicles), the GUV-SLB separation gradually changes to about 11 nm and 13 nm, with no change of roughness or uniformity of the contact area. This is consistent with no self-aggregation of the long and short DNA-linkers despite the 9 nm difference in the length of the bridges, as estimated from adhesion of vesicles with only one DNA-linker type.

To understand these findings we simulate the growth of domains with multiple binding pairs, we build on the recently developed Monte Carlo scheme for modeling adhesion by ligand-receptor pairs ^13,23^. In short, both the GUV and the SLB surfaces are discretized and decorated with DNA constructs, each occupying a single site. This approximation may result in the overestimated density of intra-membrane constructs relative to the intermembrane bridges in the equilibrium state, due to entropic considerations that have not been accounted for at this stage. However, this approximation should not affect the overall organization of bridges within adhesion domains themselves.

The system evolves in time by alternate execution of diffusion and stochastic binding/unbinding steps. The length of the time step Δ*t* is set as Δ*t* = *a*^2^*/*4*D*_DNA_, where *D*_DNA_ is the diffusion coefficient of the DNA constructs which is taken to be the same for all DNA lengths because they all have the same membrane attachments. Both the GUV and the SLB membranes are maintained on constant chemical potential ^23^, to account to for the exchange with the surrounding buffer.

In simulations, furthermore, intermittent bridges between the two membranes are formed and broken following a coarsegrained kinetics ^21,24^. DNA linkers can switch from an intra- to an inter-membrane configuration only if the opposing lattice site is empty, while breaking the bridge places the intra-membrane DNA construct on the SLB or the vesicle with equal probability. Most parameters of the simulations (Table 1) have been obtained from calibration experiments on non-functionalized membranes and SLB ^25^, and the experiments containing only one single DNA construct type ^21^. In order to simulate the experimental conditions where the contact zone expands as the DNA constructs change their conformation and bridge the two membranes, we slowly but continuously increase the contact zone, such that the concentration of bridges is always in equilibrium with the instantaneous area of contact between the two membranes ^25^.

**Table 1.**
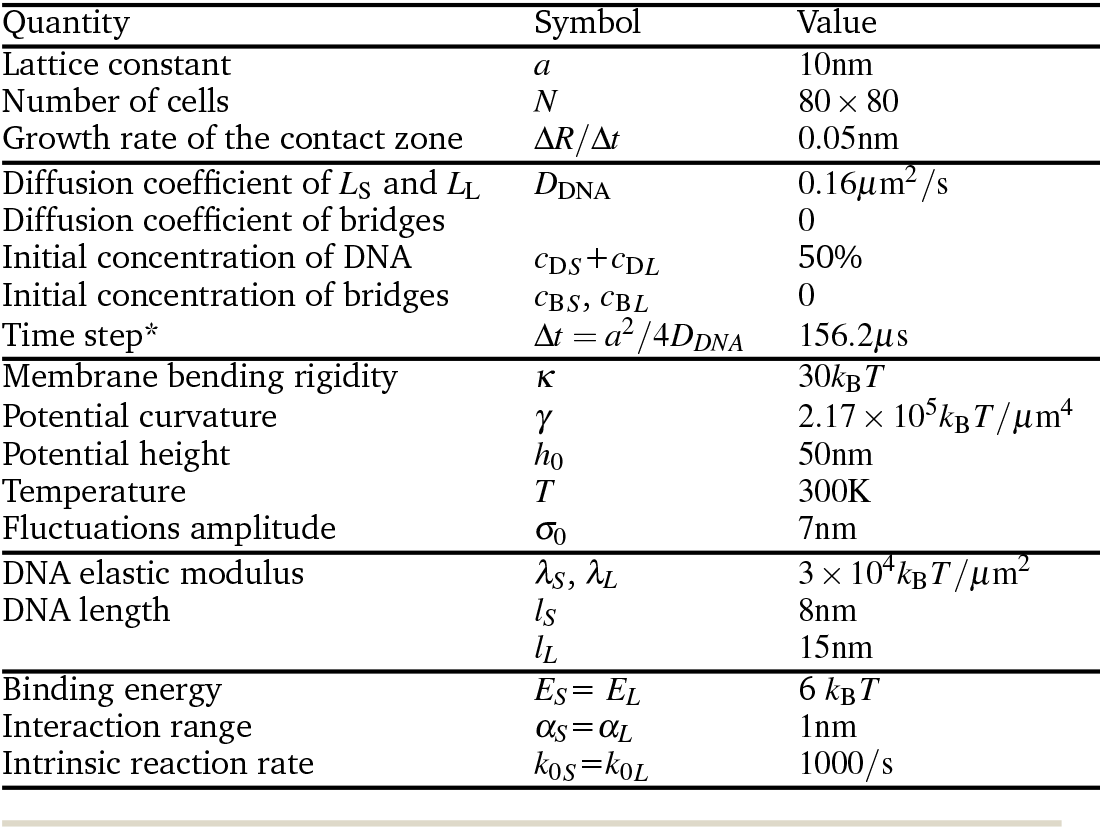
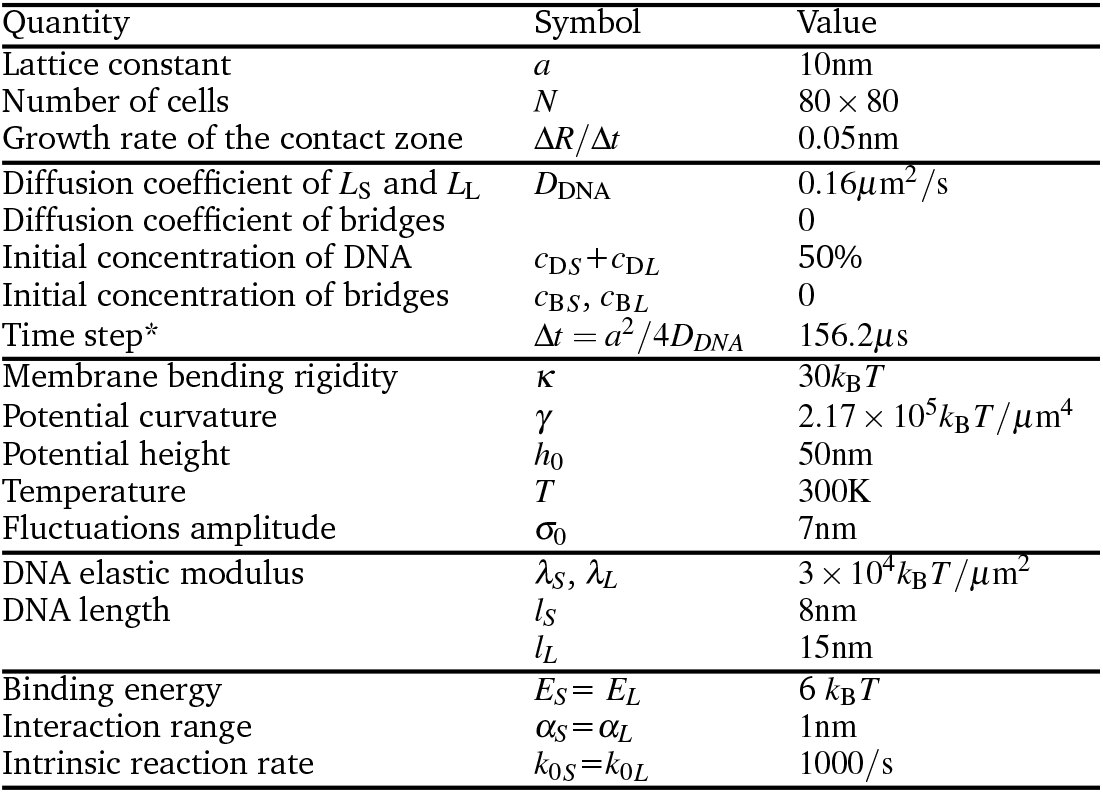
Symbols and values of parameters used in simulations with DNA linkers.

| Quantity | Symbol | Value |
| --- | --- | --- |
| Lattice constant | $a$ | 10nm |
| Number of cells | $N$ | $80 \times 80$ |
| Growth rate of the contact zone | $\Delta R/\Delta t$ | 0.05nm |
| Diffusion coefficient of $L_S$ and $L_L$ | $D_{DNA}$ | $0.16 \mu m^2/s$ |
| Diffusion coefficient of bridges |  | 0 |
| Initial concentration of DNA | $c_{DS} + c_{DL}$ | 50% |
| Initial concentration of bridges | $c_{BS}, c_{BL}$ | 0 |
| Time step* | $\Delta t = a^2/4D_{DNA}$ | 156.2μs |
| Membrane bending rigidity | $\kappa$ | $30 k_B T$ |
| Potential curvature | $\gamma$ | $2.17 \times 10^5 k_B T / \mu m^4$ |
| Potential height | $h_0$ | 50nm |
| Temperature | $T$ | 300K |
| Fluctuations amplitude | $\sigma_0$ | 7nm |
| DNA elastic modulus | $\lambda_S, \lambda_L$ | $3 \times 10^4 k_B T / \mu m^2$ |
| DNA length | $l_S$ | 8nm |
| | $l_L$ | 15nm |
| Binding energy | $E_S = E_L$ | $6 k_B T$ |
| Interaction range | $\alpha_S = \alpha_L$ | 1nm |
| Intrinsic reaction rate | $k_{0S} = k_{0L}$ | 1000/s |

A series of simulations were performed with varying relative concentrations of two types of DNA constructs in a mixture while the total number of linkers and all other parameters were kept fixed (Fig. 3). Similarly to experimental observations, we find that both short and long linkers alone develop an adhesion domain covering the entire contact zone where the bridges are uniformly distributed (Fig. 3A), in agreement with previous findings ^21^. Systematically changing the relative concentrations of the binary mixture does not macroscopically affect the nature of adhesion domains which systematically show mixing of the two bridge types, despite their relative stiffness or length .

**Fig. 3.**
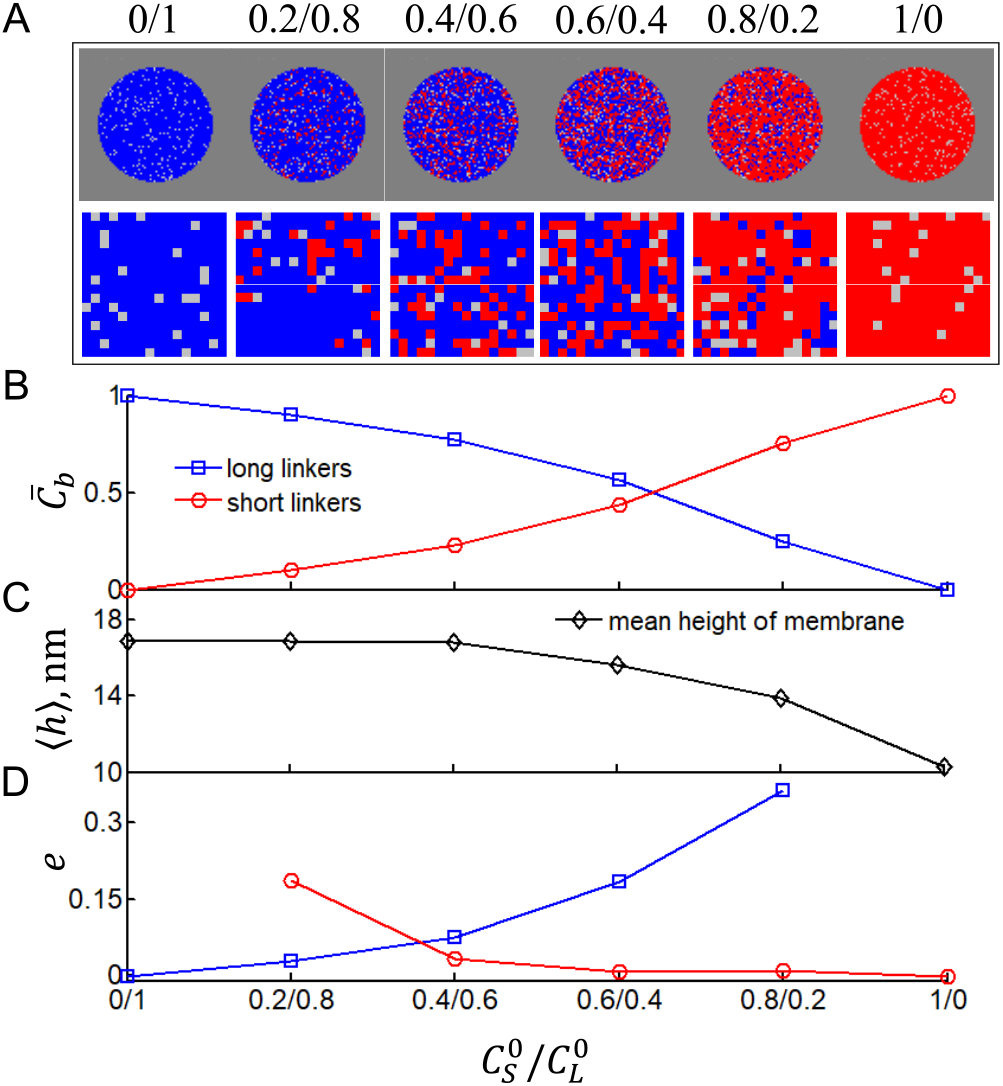
(A) Images of the (upper) contact zone and (lower) respective local domain at *t* = 7812s from two series of simulations with two types of DNA linkers. Shorter bridges are displayed in red and longer bridges in blue, free linkers are not shown. Initial relative amounts of two types of free linkers inside contact zone (red/blue) are displayed on horizontal axes. (B) Concentration of long and short bridges, 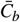, in the contact zone. (C) Average separation distance between the GUV and the SLB membrane, < h >, and (D) The excess coordination number, *e*, as a function of the binary mixture composition, showing the propensity of bridges to selfsegregate.

By increasing the fraction of short DNA linkers, the presence of short bridges in the contact zone also becomes dominant (Fig. 3B). This is naturally reflected in the average height of the GUV membrane in the contact zone, which gradually changes from 8 nm to 15 nm (Fig. 3C). However, as both linkers have about the same enthalpy of insertion, and because the energetic penalty for deforming the membrane is larger for short bridges, the equipartition of the two bridges is shifted toward mixtures with smaller concentration of long linkers, which are effectively more stable.

Notably, at the lateral resolution of about 100 nm, the roughness of the domains is similar in mixed and pure systems, consistently with the experimental finding. This gradual change in height and no variation of roughness is fully consistent with experimental findings confirming the actual nature of the domains. Even though averaged over the optical resolution, the domains appear fully mixed (Fig. 2), more detailed analysis of the simulations show that on the length scale of one DNA linker, the immediate neighborhoods of a bridge is more likely populated by the bridges of the same type (Fig. 3D). This is evident from the excess coordination number which evaluates the probability to find a nearest neighbor of the same type compared to a random distribution of bridges at the equivalent density. Obviously, pure phases show no excess coordination, consistent with the lattice gas distribution of bridges, as seen in the snapshots. However, the minority phase tends to agglomerate on very short length scales due to the membrane elasticity. As can be seen from Fig. 3E, the size of the nanodomains depends on the composition of the binary mixture, however, the effect is particularly acute when the density of a minority phase is small. Nonetheless, these nanodomains are well-below the resolution of the RICM and therefore could not be observed in our experimental systems.

In conclusions, in a joint experimental and theoretical study, we present a mimetic system showing adhesion domains containing two bridge types that are fully mixed on the length scales of optical resolution despite a 9 nm difference in their length. Using simulations, we find t hat d emixing t akes p lace o n l ength scales of few neighbors. As the direct forces between two bridges are most likely dominated by steric repulsion, the demixing on short length scales must be membrane-mediated. Namely the elastic coupling provided by the GUV bilayer is the only source of attractive, albeit weak lateral attractions. These results demonstrate that despite a significant difference length, no structural domains of one bridge type are found, presumably because both linkers independently form a lattice gas of bridges within the contact area, and an even larger difference in height would be necessary to introduce macroscopic domains with segregated bridges of one type.

## Acknowledgements

We thank the joint German Science Foundation and the French National Research Agency project SM 289/8-1, AOBJ: 652939. L. Li was further supported by the China Scholarship Council (CSC, File No. 201806185038). We are grateful to Josip Vlajčevič for the help in developing the code and performing preliminary simulations.

## Notes and references

1 E. Sackmann and A.-S. Smith, Soft Matter, 2014, 10, 1644.

2 R. Alon and K. Ley, Curr. Opin. Cell Biol., 2008, 20, 525–532.

3 M. Huse, Nat. Rev. Immunol., 2017, 17, 679.

4 C. H. Lee, H. H. Zhang, S. P. Singh, L. Koo, J. Kabat, H. Tsang, T. P. Singh and J. M. Farber, Elife, 2018, 7, e32532.

5 Y. Kaizuka, A. D. Douglass, R. Varma, M. L. Dustin and R. D. Vale, Proc. Natl. Acad. Sci. U.S.A., 2007, 104, 20296–20301.

6 S. F. Banani, H. O. Lee, A. A. Hyman and M. K. Rosen, Nat. Rev. Mol. Cell Biol., 2017, 18, 285–298.

7 T. R. Weikl, M. Asfaw, H. Krobath, B. Różycki and R. Lipowsky, Soft Matter, 2009, 5, 3213.

8 E. M. Schmid, M. H. Bakalar, K. Choudhuri, J. Weichsel, H. S. Ann, P. L. Geissler, M. L. Dustin and D. A. Fletcher, Nat. Phys., 2016, 12, 704–711.

9 S. F. Fenz, T. Bihr, D. Schmidt, R. Merkel, U. Seifert, K. Sengupta and A.-S. Smith, Nat. Phys., 2017, 13, 906–913.

10 M. Weiss, J. P. Frohnmayer, L. T. Benk, B. Haller, J.-W. Janiesch, T. Heitkamp, M. Börsch, R. B. Lira, R. Dimova, R. Lipowsky et al., Nat. Mater., 2018, 17, 89.

11 T. R. Weikl and R. Lipowsky, Biophys. J., 2004, 87, 3665–3678.

12 P. K. Tsourkas, M. L. Longo and S. Raychaudhuri, Biophys. J., 2008, 95, 1118–1125.

13 J. Vlajčevič and A.-S. Smith, In preparation. Paper on experiments with DNA-linkers done at CINaM, CNRS, Marseille.

14 M. Chung, B. J. Koo and S. G. Boxer, Faraday Discuss., 2013, 161, 333–345.

15 S. F. Shimobayashi, B. M. Mognetti, L. Parolini, D. Orsi, P. Cicuta and L. D. Michele, Phys. Chem. Chem. Phys., 2015, 17, 15615–15628.

16 S. J. Bachmann, J. Kotar, L. Parolini, A. Šarič, P. Cicuta, L. D. Michele and B. M. Mognetti, Soft Matter, 2016, 12, 7804–7817.

17 L. Parolini, B. M. Mognetti, J. Kotar, E. Eiser, P. Cicuta and L. Di Michele, Nat. Commun., 2015, 6, 5948.

18 O. A. Amjad, B. M. Mognetti, P. Cicuta and L. Di Michele, Langmuir, 2017, 33, 1139–1146.

19 W. T. Kaufhold, R. A. Brady, J. M. Tuffnell, P. Cicuta and L. Di Michele, Bioconjug. Chem., 2019.

20 B. M. Mognetti, P. Cicuta and L. Di Michele, Rep. Prog. Phys., 2019, 82, 116601.

21 M. A. Kamal, K. Sengupta, F. Thibaudau, J. Vlajčevič and A.-S. Smith, In preparation. Paper on experiments with DNA-linkers done at CINaM, CNRS, Marseille.

22 L. K. Tamm and H. M. McConnell, Biophys. J., 1985, 47, 105–113.

23 T. Bihr, U. Seifert and A.-S. Smith, New J. Phys., 2015, 17, 083016.

24 T. Bihr, U. Seifert and A.-S. Smith, Phys. Rev. Lett., 2012, 109, 258101.

25 D. Schmidt, T. Bihr, S. Fenz, R. Merkel, U. Seifert, K. Sengupta and A.-S. Smith, BBA-Mol. Cell Res., 2015, 1853, 2984–2991.

